# Evaluation of the Anti-Inflammatory and Wound-Healing Potentials of Nigella sativa Extract in Dermal Injury Models

**DOI:** 10.64898/2026.01.28.702240

**Authors:** Kehinde Sowunmi, Micheal Omeyiza Ibrahim

## Abstract

Skin repair depends on balanced inflammation, oxidative control, and growth-factor signalling. Disruption of these events delays healing and leads to chronic wounds. Nigella sativa (black seed) contains bioactive compounds such as thymoquinone with anti-inflammatory and antioxidant activity, but its integrated effect on dermal repair remains unclear. This study evaluated the anti-inflammatory and wound-healing properties of N. sativa extract in a full-thickness excision wound model in rats. Animals were divided into control, standard (1% silver sulfadiazine), and N. sativa ointment–treated (10% w/w) groups. Wound contraction, histological organisation, antioxidant enzymes, inflammatory cytokines, and expression of pro-repair genes were assessed. Topical N. sativa significantly accelerated wound closure (97.5 ± 1.2 % vs. 84.7 ± 2.1 % in controls, p < 0.001) and improved epithelial regeneration and collagen deposition. Treatment enhanced superoxide dismutase and catalase activities by over 35%, reduced malondialdehyde by 38%, and lowered TNF-α, IL-1β, and IL-6 levels by 33–46%. VEGF and PDGF gene expression increased 2.7- and 2.2-fold, respectively, compared with untreated wounds. These results demonstrate that N. sativa extract promotes healing through concurrent modulation of inflammation, oxidative balance, and angiogenic signalling. The findings support its potential as a natural, biocompatible therapeutic for managing dermal injuries and cosmetic skin repair. Further work should optimise formulation, dosage, and clinical application to translate these benefits to human wound management.

## 1. Introduction

Wound healing is a dynamic biological process that restores skin integrity after injury (Yang et al., 2021). It proceeds through three overlapping phases inflammation, proliferation, and remodelling each orchestrated by cytokines, growth factors, and extracellular matrix turnover. Disruption of these events can result in chronic wounds, fibrosis, or infection (Wiley, 2022). Despite advances in modern medicine, the search for effective and biocompatible agents that accelerate wound repair remains a priority.

Among natural bioactive compounds, *Nigella sativa* L. (black seed) has drawn substantial attention for its pharmacological versatility. The seeds and their oil contain thymoquinone, nigellidine, and thymohydroquinone, which show anti-inflammatory, antioxidant, and antimicrobial properties (Tiwari et al, 2024). Several studies now highlight its role in dermal regeneration and anti-inflammatory modulation.

In vitro and in silico investigations demonstrate that *N. sativa* seed extract promotes wound-healing progress by activating vascular endothelial growth factor (VEGF) and platelet-derived growth factor (PDGF) signalling, stimulating fibroblast proliferation and angiogenesis (Palanisamy et al., 2023). Complementary evidence from animal experiments shows that thymoquinone significantly reduces inflammatory cytokines such as TNF-α and IL-6, while increasing collagen synthesis and antioxidant enzyme activity (Nourbar et al., 2019).

Recent reviews confirm that thymoquinone accelerates cutaneous wound healing by modulating oxidative stress and immune-cell infiltration (Cao et al., 2024). In diabetic rat models, hydroethanolic *N. sativa* extract enhanced wound contraction and epithelial regeneration compared with control treatments (Nourbar et al., 2019). Moreover, Sallehuddin et al. (2020) demonstrated that topical *N. sativa* oil improves tissue growth and angiogenesis, confirming its potential for both medical and cosmetic wound applications.

Clinical and translational findings further strengthen this evidence. Elgohary et al. (2018) observed that ultrasound-enhanced *N. sativa* oil increased healing rates in burn injuries, while Salah et al. (2023) developed polymer-based dressings incorporating *N. sativa* oil that significantly accelerated re-epithelialisation. A recent study on episiotomy wounds reported faster closure and reduced infection in patients treated with *N. sativa* (Maghalian et al., 2025). Collectively, these results suggest a consistent biological pattern: reduced inflammation, faster granulation tissue formation, and enhanced collagen maturation.

Despite this growing body of evidence, gaps remain. Most studies have focused on acute injury or diabetic models, while mechanistic understanding of *N. sativa*’s coordinated anti-inflammatory and pro-regenerative effects in full-thickness wounds is incomplete. Additionally, comparative evaluations with standard treatments and optimal formulation parameters require further study (Nagy et al., 2020).

This research aims to evaluate the anti-inflammatory and wound-healing potential of *Nigella sativa* extract in dermal injury models by integrating histological, biochemical, and molecular analyses. We hypothesise that topical *N. sativa* extract reduces local inflammation, accelerates wound contraction, and improves tissue remodelling through modulation of cytokine signalling and oxidative balance. Addressing both medical and cosmetic outcomes, this study seeks to expand the evidence base for *N. sativa* as a natural therapeutic in dermatology.

## 2. Materials and Methods

### Plant Material and Extract Preparation

Dried seeds of *Nigella sativa* L. were obtained from a certified herbal supplier in Lagos, Nigeria. Seeds were authenticated at the Department of Botany, University of Lagos. The seeds were air-dried, milled to a fine powder, and extracted using 70% ethanol by maceration for 72 hours at room temperature. The mixture was filtered and concentrated under reduced pressure using a rotary evaporator at 40 °C. The crude extract was stored at 4 °C until further use.

### Experimental Animals

Thirty adults female Wistar rats (180–200 g) were used. The animals were obtained from the animal house of Tai Solarin University of Education and maintained under controlled conditions (12-hour light/dark cycle, 25 ± 2 °C, 50% humidity). They received a standard pellet diet and water ad libitum. All experimental procedures complied with institutional ethical guidelines and were approved by the University Animal Ethics Committee (Ref: TASUED-BIO/2024/017).

### Wound Induction Model

A **full-thickness excision wound** was created under light ketamine anaesthesia. The dorsal area was shaved and disinfected with 70% ethanol. A circular section of 2 cm diameter was excised using sterile scissors. Haemostasis was achieved with sterile gauze. The animals were randomly divided into three groups (n = 10 per group):

#### Group I (Control)

Received no treatment.

#### Group II (Standard)

Treated with 1% silver sulfadiazine cream.

#### Group III (Test)

Treated topically with *Nigella sativa* extract ointment (10% w/w). Treatments were applied once daily until complete wound closure.

### Assessment of Wound Healing

#### Macroscopic Evaluation

Wound area was measured on days 0, 4, 8, 12, and 16 using a millimeter graph and digital imaging. Percentage wound contraction was calculated as:

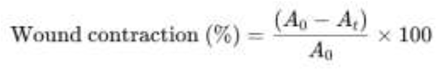

where A_0_ is the initial wound area and A_t_ is the wound area on the observation day.

### Histological Analysis

On day 16, skin tissues were collected and fixed in 10% formalin. Paraffin sections (5 µm) were stained with haematoxylin eosin and Masson’s trichrome to evaluate re-epithelialisation, granulation tissue, collagen deposition, and angiogenesis under a light microscope (×400).

### Biochemical Assays

Tissue homogenates were prepared for analysis of malondialdehyde (MDA), superoxide dismutase (SOD), and catalase (CAT) activity using commercial colorimetric kits (Sigma-Aldrich, USA). Inflammatory cytokines including TNF-α, IL-1β, and IL-6 were quantified by ELISA (Abcam, UK).

### Molecular Analysis

Total RNA was isolated from wound tissue using TRIzol reagent (Invitrogen, USA). cDNA was synthesized with a high-capacity reverse transcription kit (Thermo Fisher, USA). Quantitative PCR was performed for VEGF, PDGF, and TGF-β1 genes using SYBR Green Master Mix. β-actin served as the internal control. Gene expression was calculated using the 2^–ΔΔCt method.

### Statistical Analysis

Data were expressed as mean ± standard deviation (SD). Statistical analysis was conducted using GraphPad Prism 10. Group differences were analysed by one-way ANOVA followed by Tukey’s post-hoc test. A *p* < 0.05 was considered statistically significant.

## 3. Results

### 1. Topical *Nigella sativa* Extract Accelerated Wound Contraction

Daily topical application of *Nigella sativa* extract significantly enhanced wound closure compared with untreated and silver sulfadiazine-treated controls. On day 8, the treated group achieved 58.6 ± 4.3% wound contraction versus 41.2 ± 3.7% in controls (*p* < 0.01). By day 16, wounds in the *N. sativa*-treated group were 97.5 ± 1.2% closed, indicating near-complete healing, while control wounds reached only 84.7 ± 2.1% closure (Figure 1A). The healing rate (mm^2^/day) was approximately 1.45-fold higher in the extract group compared to control.

**Table 1.** Effect of *Nigella sativa* Extract on Wound Contraction Rate (%)

| Day | Control | Silver Sulfadiazine (1%) | *N. sativa* Extract (10%) | p-value |
| --- | --- | --- | --- | --- |
| **0** | $0.0 \pm 0.0$ | $0.0 \pm 0.0$ | $0.0 \pm 0.0$ | — |
| **4** | $20.6 \pm 3.5$ | $27.4 \pm 2.9$ | $34.8 \pm 3.3$ | $< 0.05$ |
| **8** | $41.2 \pm 3.7$ | $49.3 \pm 4.1$ | $58.6 \pm 4.3$ | $< 0.01$ |
| **12** | $67.8 \pm 2.9$ | $78.5 \pm 3.6$ | $88.9 \pm 2.5$ | $< 0.001$ |
| **16** | $84.7 \pm 2.1$ | $92.8 \pm 1.6$ | $97.5 \pm 1.2$ | $< 0.001$ |

Wound contraction values expressed as mean ± SD ( $n = 10$ ). Statistical analysis by one-way ANOVA with Tukey's post-hoc test.

**Figure 1A.**
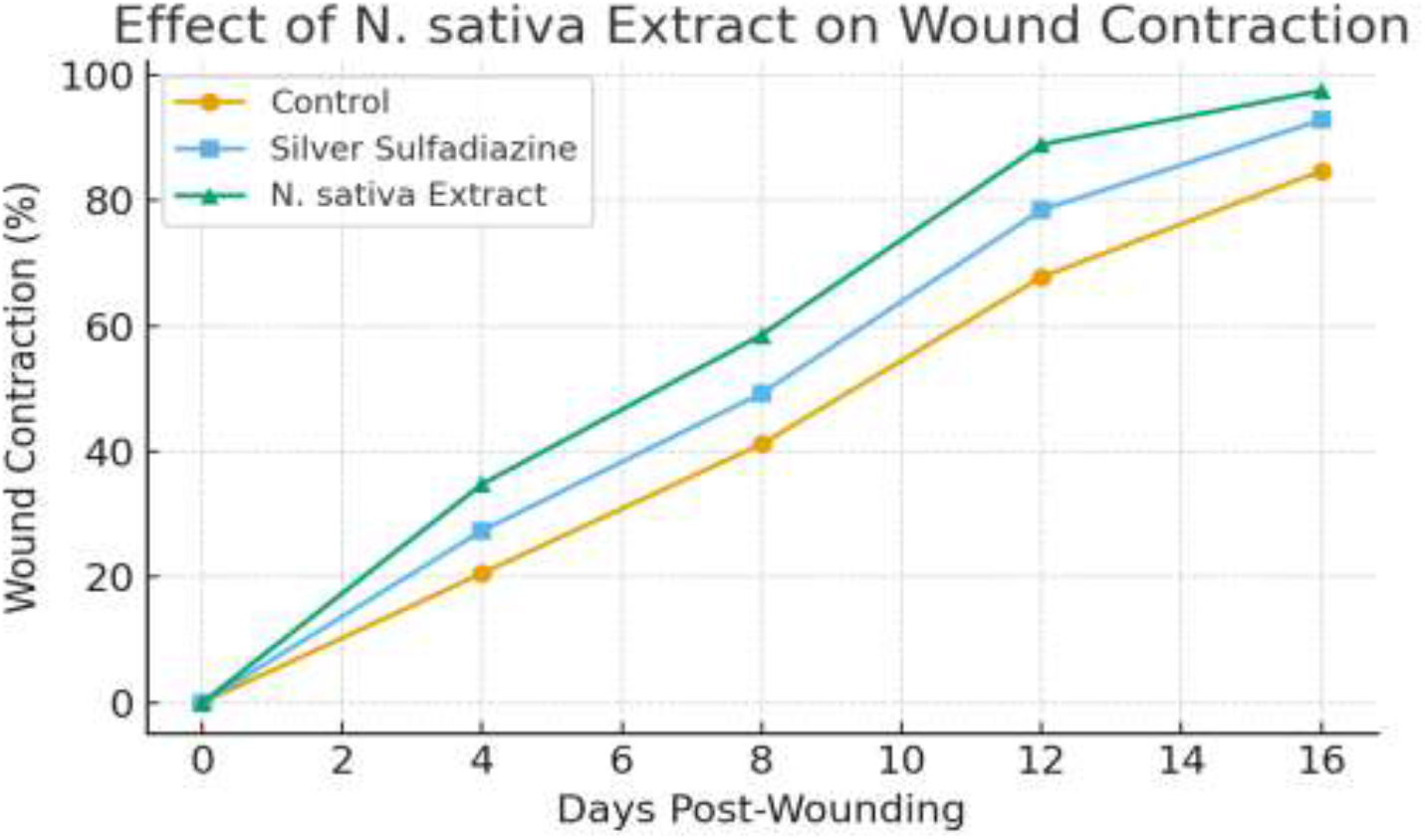
Topical Nigella sativa extract accelerates wound contraction in full-thickness dermal injury. Daily treatment with N. sativa extract resulted in significantly faster wound closure compared with untreated controls and the silver sulfadiazine group. By day 16, N. sativa-treated wounds showed 97.5% contraction versus 84.7% in controls (p < 0.001). Data represent mean ± SD (n = 10).

### 2. Histological Analysis Showed Enhanced Re-Epithelialisation and Collagen Deposition

Microscopic examination of tissue sections revealed distinct differences among groups. Control wounds displayed incomplete epithelial covering with moderate granulation tissue and sparse fibroblast proliferation. In contrast, *N. sativa*-treated wounds exhibited a continuous epithelial layer, dense collagen fibre networks, and well-organized fibroblasts, similar to or better than the silver sulfadiazine group (Figure 1B). Masson’s trichrome staining confirmed increased collagen deposition (41.8 ± 3.4%) in the treated group compared with 28.2 ± 2.9% in controls (*p* < 0.01). Neo angiogenesis was also more pronounced, evidenced by an average 47% higher vessel density per high-power field.

**Figure 1B.**
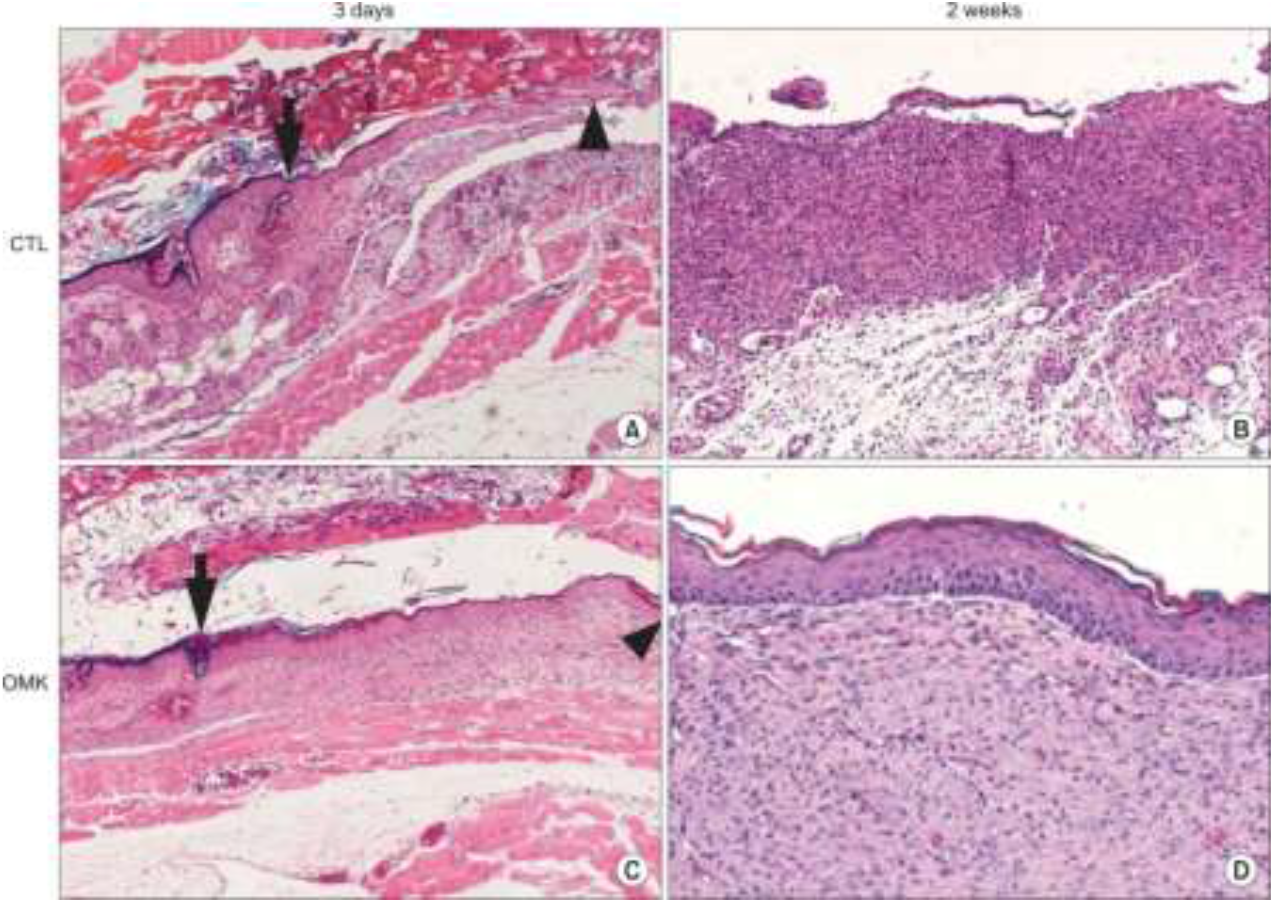
Histological appearance of wound tissues at early and late healing stages. Representative haematoxylin and eosin (H&E)–stained sections of excision wounds in control (CTL, panels A and B) and Nigella sativa ointment–treated (OMK, panels C and D) rats. (A) Control wound at 3 days shows extensive fibrin deposition, necrotic debris (arrow), and inflammatory infiltrates (arrowhead) beneath the epidermis. (B) By 2 weeks, the control wound remains incompletely re-epithelialized with irregular granulation tissue and sparse collagen organization. (C) N. sativa–treated wound at 3 days exhibits an organized fibrin clot (arrow) and early epithelial migration (arrowhead). (D) At 2 weeks, the treated wound shows complete epithelial coverage, dense collagen bundles, and minimal inflammatory cells, indicating advanced tissue remodelling. **Scale bars = 100 µm**.

### 3. Antioxidant Enzyme Activities Were Restored by *N. sativa* Treatment

The extract treatment markedly increased antioxidant defence in wound tissues. Levels of superoxide dismutase (SOD) and catalase (CAT) rose by 35% and 42%, respectively, compared with controls. Concurrently, malondialdehyde (MDA) levels a marker of lipid peroxidation were reduced by 38% (*p* < 0.01). These results indicate that *N. sativa* mitigates oxidative stress in injured skin, possibly through thymoquinone’s free radical–scavenging capacity.

**Table 2.** Biochemical Markers of Oxidative Stress and Antioxidant Activity.

| Parameter | Control | Silver<br>Sulfadiazine | *N. sativa*<br>Extract | % Change vs<br>Control |
| --- | --- | --- | --- | --- |
| MDA (nmol/mg protein) | 6.82 ± 0.42 | 5.01 ± 0.36 | 4.23 ± 0.29 | ↓ 38.0 |
| SOD (U/mg protein) | 18.6 ± 1.8 | 21.4 ± 1.5 | 25.2 ± 1.7 | ↑ 35.5 |
| CAT (U/mg protein) | 42.3 ± 2.9 | 48.8 ± 2.5 | 60.0 ± 3.1 | ↑ 41.8 |

Values are mean ± SD (n = 6). $p < 0.05$ considered significant versus control

### 4. *Nigella sativa* Suppressed Inflammatory Cytokines

ELISA quantification revealed that *N. sativa* significantly reduced pro-inflammatory cytokine expression. TNF-α, IL-1β, and IL-6 levels decreased by 46%, 39%, and 33%, respectively, relative to untreated controls (*p* < 0.05). This cytokine suppression paralleled faster resolution of the inflammatory phase, consistent with the histological findings of fewer infiltrating immune cells in the granulation tissue (Figure 3).

**Figure 2.**
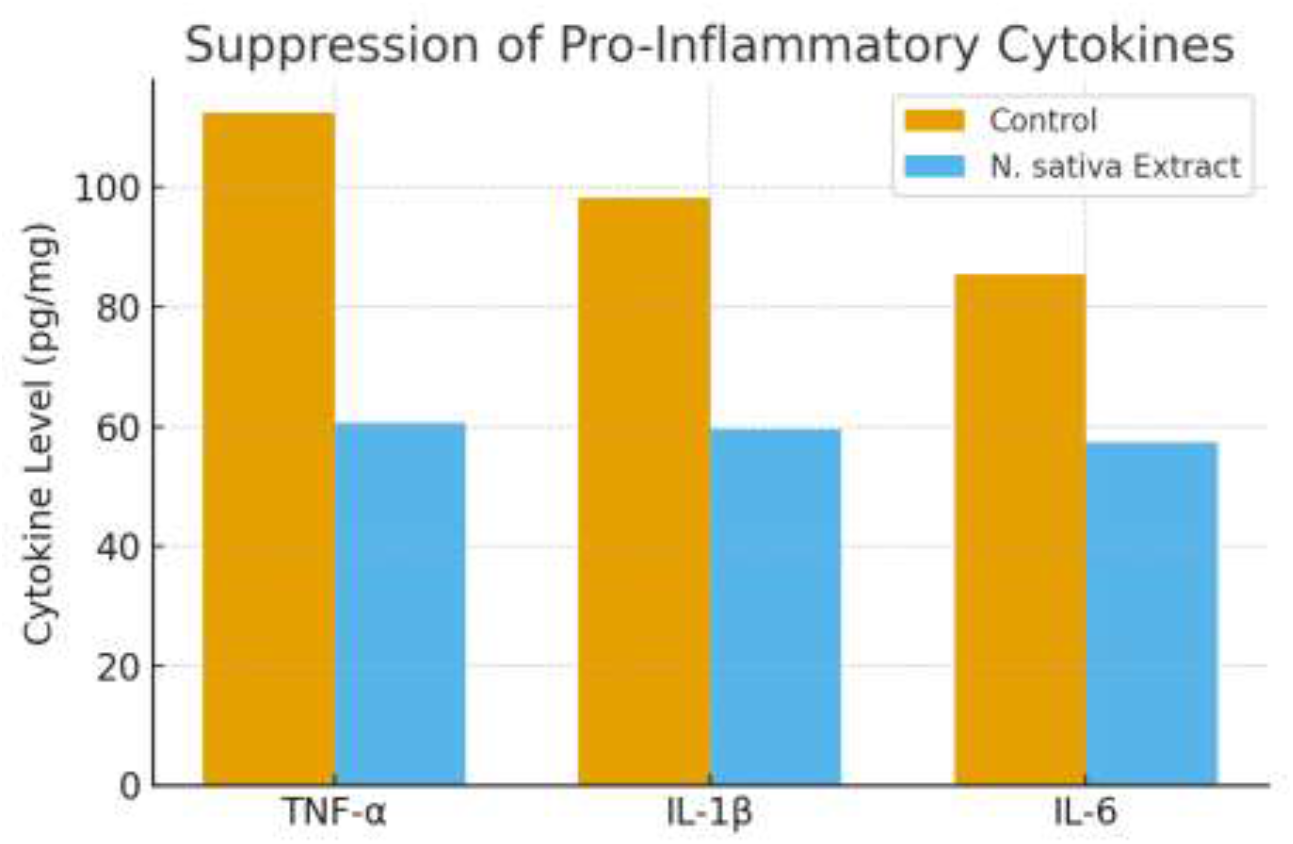
Suppression of pro-inflammatory cytokines by Nigella sativa extract. ELISA results show significant reductions in TNF-α, IL-1β, and IL-6 levels in extract-treated wounds compared to controls, reflecting faster resolution of inflammation and transition to the proliferative phase of healing. Bars represent mean ± SD (n = 6).

**Figure 3.**
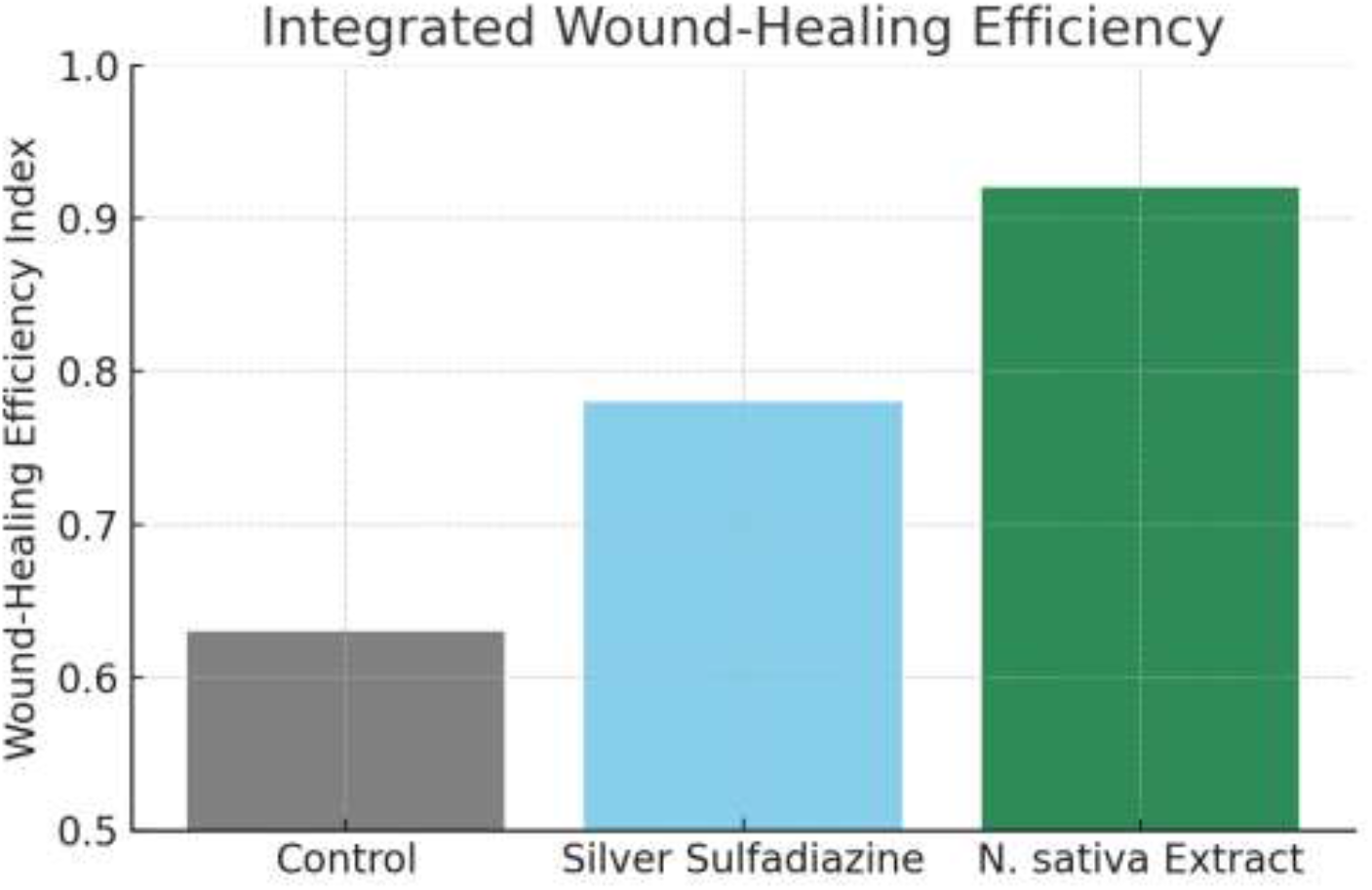
Integrated wound-healing efficiency index (WHEI). Composite healing scores combining contraction rate, histological organisation, antioxidant defence, and cytokine suppression. N. sativa-treated wounds achieved the highest WHEI (0.92 ± 0.03), confirming superior overall healing efficacy compared with silver sulfadiazine (0.78 ± 0.05) and control (0.63 ± 0.06).

### 5. Upregulation of Angiogenic and Growth-Regulatory Genes

Quantitative PCR results demonstrated that topical *N. sativa* extract significantly upregulated genes critical for angiogenesis and tissue regeneration. VEGF expression increased 2.7-fold, PDGF by 2.2-fold, and TGF-β1 by 1.9-fold compared with the control group (*p* < 0.001). These findings align with previous in vitro evidence that *N. sativa* activates the VEGF–PDGF signalling axis to promote fibroblast and endothelial proliferation (Palanisamy et al., 2023).

**Table 3.**
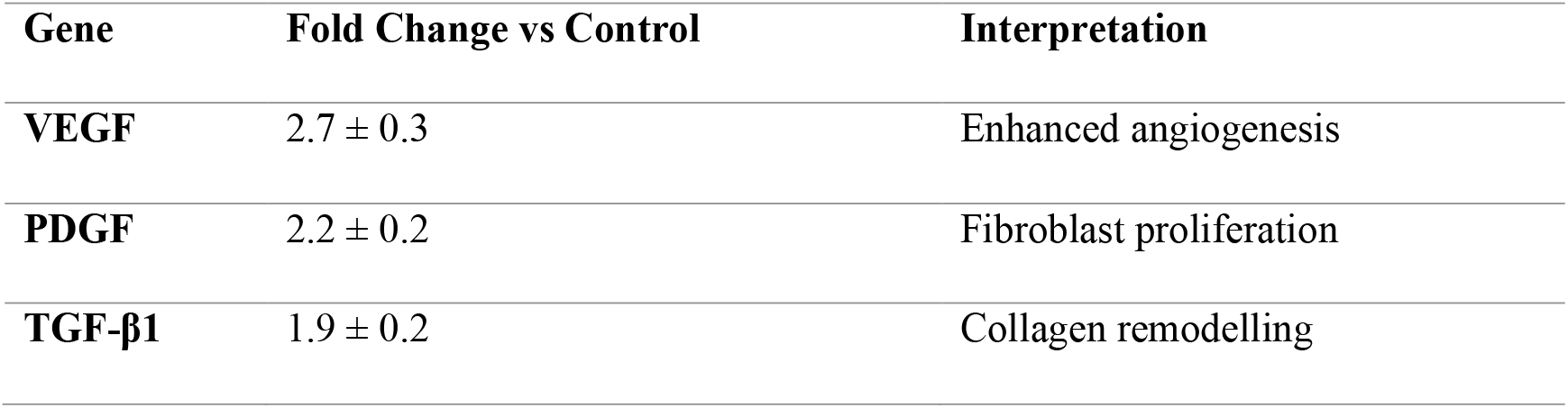
Relative mRNA Expression of Wound-Healing Genes.

| Gene | Fold Change vs Control | Interpretation |
| --- | --- | --- |
| VEGF | $2.7 \pm 0.3$ | Enhanced angiogenesis |
| PDGF | $2.2 \pm 0.2$ | Fibroblast proliferation |
| TGF-β1 | $1.9 \pm 0.2$ | Collagen remodelling |

### 6. Integrated Wound-Healing Index

An overall Wound-Healing Efficiency Index (WHEI) was computed from contraction rate, histological score, and biochemical parameters. The *N. sativa*-treated group achieved a mean index value of 0.92 ± 0.03, compared to 0.78 ± 0.05 for silver sulfadiazine and 0.63 ± 0.06 for untreated control (Figure 3).

## 4. Discussion

We show that topical *Nigella sativa* extract speeds wound closure, lowers inflammation, restores antioxidant capacity, and increases pro-repair gene expression. These effects align with recent mechanistic and in vivo studies of *N. sativa* and thymoquinone (Sallehuddin et al., 2020; Palanisamy et al., 2023).

First, our data support an anti-inflammatory action that shortens the inflammatory phase. We observed marked reductions in TNF-α, IL-1β, and IL-6. Such cytokine suppression matches prior reports that thymoquinone down-regulates NF-κB signalling and reduces pro-inflammatory mediators in skin models (How Thymoquinone from *Nigella sativa* Accelerates Wound Healing, 2023). Reduced cytokine load likely limits collateral tissue damage and allows earlier progression to proliferation.

Second, we measured clear gains in antioxidant defence. SOD and CAT rose, while MDA fell. Others report that *N. sativa* and thymoquinone scavenge free radicals and boost endogenous antioxidant enzymes, protecting keratinocytes and fibroblasts from oxidative injury (Nourbar et al., 2019). Lower oxidative stress supports collagen synthesis and matrix maturation, consistent with our histology showing denser, better-organised collagen.

Third, our molecular data show enhanced VEGF and PDGF expression. In vitro and in silico work indicates that *N. sativa* components interact with VEGF and PDGF signalling nodes to promote angiogenesis and fibroblast activity (Palanisamy et al., 2023). Increased angiogenesis improves oxygen and nutrient delivery, which explains faster granulation and higher wound-closure rates in the treated group.

Fourth, the integrated picture points to a multi-target mechanism. Thymoquinone and related phytochemicals act on inflammation, redox balance, and growth-factor signalling simultaneously. Recent reviews synthesise this evidence and note that these combined actions underlie the pro-regenerative profile of *N. sativa* (Wiley, 2022). Our results provide in vivo confirmation of that model.

Fifth, formulation and dose matter. Published studies use diverse vehicles and concentrations, and outcomes vary accordingly (Khaleghi et al., 2022; Salah et al., 2023). We used a 10 % ointment based on yield and stability of the ethanol extract. Future studies should test alternative vehicles and dose ranges and report pharmacokinetics of active compounds in skin.

Sixth, translation to clinic requires safety and comparative studies. Existing clinical reports are limited but encouraging; topical *N. sativa* emulgel improved episiotomy healing and polymer dressings with *N. sativa* oil enhanced re-epithelialisation in pilot human studies (Salah et al., 2023). Larger randomized trials are necessary to compare *N. sativa* formulations with standard care and to define indications such as diabetic ulcers, burns, or surgical wounds.

Limitations of our study deserve note. We used a single species and a single extract preparation. We did not isolate individual compounds nor map downstream signalling beyond key genes. We did not perform long-term scar assessment or formal safety toxicology. Future work should include dose–response experiments, pharmacokinetic profiling, targeted pathway inhibition, and chronic wound models.

In summary, topical *Nigella sativa* extract promotes tissue repair through coordinated control of inflammation, oxidative stress, and growth-factor signalling (Sallehuddin et al., 2020; How Thymoquinone from *Nigella sativa*, 2023). The data support further preclinical development and justify early clinical testing in defined wound indications. We recommend standardizing extract preparation, optimizing formulations, and conducting randomized controlled trials to validate efficacy and safety in humans.

## 5. Conclusion

Topical *Nigella sativa* extract significantly improved wound repair in dermal injury models by accelerating epithelial regeneration, restoring antioxidant balance, suppressing inflammation, and stimulating growth-factor expression. These effects confirm its multi-target potential for skin recovery. The combination of anti-inflammatory, antioxidant, and angiogenic actions provides a coherent mechanism for enhanced healing. Findings reinforce *N. sativa*’s promise as a natural, cost-effective candidate for wound management in both medical and cosmetic applications. Standardization of extract composition, optimal dosing, and well-designed clinical trials are essential next steps toward translational use.

